# Computational optimization of two-photon holographic stimulation sites *in vivo*

**DOI:** 10.1101/2025.07.31.667911

**Authors:** Marcus A. Triplett, Edgar Bäumler, Alex Prodan, Rokas Stonis, Darcy S. Peterka, Michael Häusser, Liam Paninski

## Abstract

Determining the intricate structure and function of neural circuits requires the ability to precisely manipulate circuit activity. Two-photon holographic optogenetics has emerged as a powerful tool for achieving this via flexible excitation of user-defined neural ensembles. However, the precision of two-photon optogenetics has been constrained by off-target stimulation, an effect where proximal non-target neurons can be unintentionally activated due to imperfect spatial confinement of light onto target neurons. Here, we introduce a real-time computational method to mitigating off-target stimulation that first empirically samples each neuron’s sensitivity to stimulation at proximal locations, and then optimizes stimulation sites using a fast, interpretable model based on adaptive non-negative basis function regression (NBFR). NBFR is highly scalable, completing model fitting for hundreds of neurons in just a few seconds and then optimizing stimulation sites in several hundred milliseconds per stimulus – fast enough for most closed-loop behavioral experiments. We characterize the performance of our approach in both simulations and *in vivo* experiments in mouse hippocampus, showing its efficacy under realistic experimental conditions. Our results thus establish NBFR-based photostimulus optimization as an important addition to an emerging computational toolkit for precise yet scalable holographic optogenetics.

## Introduction

Determining the specific patterns of neural activity giving rise to behaviors or cognitive functions requires precise causal interventions *in vivo*. Holographic optogenetics has emerged as a crucial technology for this purpose due to its ability to flexibly deliver two-photon excitation to user-selected ensembles of neurons with near single-cell resolution [Papagiakoumou et al., 2010, Packer et al., 2012, Prakash et al., 2012, Packer et al., 2015, Yang et al., 2018, Adesnik and Abdeladim, 2021]. When combined with calcium imaging, holographic optogenetics enables minimally-invasive “all-optical” reading and writing of neural activity patterns in real time, allowing for closed-loop probing of neural circuits [Zhang et al., 2018, Hira, 2024, Draelos et al., 2025]. However, despite the significant improvement in spatial precision over one-photon optogenetics and electrical stimulation, holographic optogenetics has still been critically limited by the problem of off-target stimulation (OTS): the inadvertent activation of nearby non-target neurons due to imperfect confinement of light onto the somas of target neurons [Adesnik and Abdeladim, 2021, Triplett et al., 2023, Lees et al., 2024]. While the use of soma-targeted opsins [Baker et al., 2016, Shemesh et al., 2017] such as ST-ChroME [Mardinly et al., 2018, Sridharan et al., 2022] and ChRmine [Marshel et al., 2019] has improved the precision of holographic optogenetics over untargeted opsins in densely expressing preparations, OTS still remains a fundamental limitation to the precision and specificity of optogenetic manipulation experiments, and has been described as an outstanding challenge for this technology [Adesnik and Abdeladim, 2021].

A computational solution to the OTS problem is highly appealing, as it would require no additional hardware, optical calibration, or further expenses. However, any such approach should satisfy two important criteria. First, the approach should operate in real time, so that an experimenter can quickly adjust stimulation patterns without significant overhead in runtime. Second, the approach should be scalable: experiments often involve reading activity from and writing activity into many hundreds of neurons, and future mesoscale systems will likely increase this number into the thousands [Abdeladim et al., 2023]. A prior approach [Bounds et al., 2023] considered optimizing the laser power and stimulation frequency to recreate specific, naturalistic activity patterns, but did not attempt to explicitly minimize off-target activation. By contrast, the first method to explore the direct minimization of OTS by simultaneously optimizing the exact stimulation sites and laser powers was introduced by ref. [Triplett et al., 2023] via the “Bayesian target optimization” algorithm. However, Bayesian target optimization was only tested in simulations derived from previously collected slice recordings (and therefore has not been tested for suitability in real-time experiments or under *in vivo* conditions), and is based on Gaussian process regression techniques, which can involve computationally intensive and slow operations on large matrices, precluding use for real-time purposes.

Here we report on the development of a computational approach to overcoming OTS based on real-time probing of neural response properties and subsequent fast optimization of holographic stimulation sites. We characterized the performance of our method in detailed simulations, and tested it in experiments in hippocampal area CA1 of awake behaving mice. We further determined computation time requirements, which demonstrated suitability for use in real-time experiments. Together, our results show that photostimulus optimization can considerably mitigate off-target activation in practice with minimal computational overhead.

## Results

### Optimizing photostimulation patterns using adaptive non-negative basis function regression

Prior work has shown that neurons may respond to two-photon optogenetic stimulation even when the stimulus is outside of the perimeter of the cell soma (~5-15 *µ*m laterally, ~20-50 *µ*m axially [Baker et al., 2016, Lees et al., 2024]). This phenomenon has been hypothesized to result from the spatial extent of the optical point spread function, together with residual expression of opsin molecules in the proximal dendrites (both within the stimulation plane as well as immediately above and below) despite the use of soma-targeted opsins. Thus, if neurons are closely packed, attempting to stimulate one neuron by focusing light directly at its soma can frequently activate neighboring neurons as well [Adesnik and Abdeladim, 2021]. Optimizing a stimulation pattern therefore requires determining the specific spatial region where a neuron can be activated. We refer to these regions as “optogenetic receptive fields” (ORFs) as described in previous work [Triplett et al., 2023].

To determine these ORFs, our approach begins by using holographic optogenetics to probe where neurons respond to stimulation in a small grid of stimulation sites surrounding each neuron (Figure 1a,b). For improved efficiency, we remove any “redundant” stimulation sites that are within several micrometers of others due to overlapping grids from neighboring neurons. Then, to accelerate ORF mapping, we deliver stimulation to multiple (typically 10) sites simultaneously to map ORFs in parallel. This also allows us to probe how neurons integrate two-photon excitation from multiple simultaneous nearby holographic targets; our hypothesis being that such data would ensure that we can optimize stimulation patterns targeting ensembles of neurons, and not just single cells.

**Figure 1:**
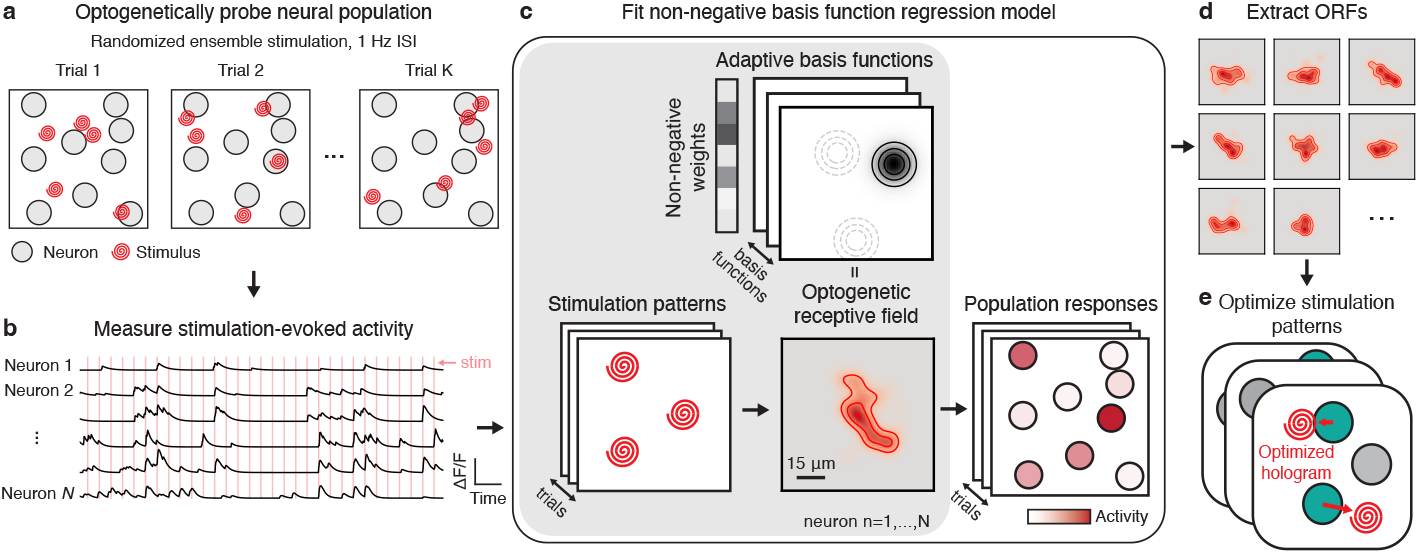
Real-time approach to mapping optogenetic response properties and optimizing stimulation. **a**, Simultaneous holographic excitation of multiple stimulation sites is used to evoke calcium transients across all neurons within a desired FOV. **b**, Stimulation-evoked neural activity is extracted and used for modeling. **c**, Adaptive non-negative basis function regression (NBFR) uses stimulation patterns and neural population responses to identify each neuron’s “optogenetic receptive field” – i.e., the fluorescence response evoked by stimulation at each nearby point in space. Solid concentric circles represent first basis function, dashed concentric circles show locations of other basis functions. **d-e** NBFR-computed ORFs (**d**) are used to optimize the exact ensemble stimulation pattern (**e**) to minimize OTS while still exciting the target neurons. In **e**, each stacked panel refers to a different stimulation pattern to be optimized. ORFs in this overview estimated from CA1 neurons.

Given the neural population responses to stimulation, we next use an adaptive non-negative basis function regression method (NBFR, see Methods) to fit statistical models describing the ORFs (Figure 1c,d). NBFR uses a non-negative mixture of Gaussians to model the spatial structure of the stimulus responses – a more tractable finite-basis analogue of the infinite-dimensional Gaussian process approach explored in our previous work [Triplett et al., 2023]. Finally, we leverage the inferred ORFs to quickly compute adjustments to ensemble stimulation patterns that optimally avoid non-target neurons while still activating target neurons (Figure 1e).

### NBFR enables learning of photostimulus response properties

We performed a series of simulations to characterize the behavior of NBFR under various experimentally-relevant con-texts. We randomly positioned neurons in an artificial field of view (FOV, Figure 2a), and assigned each neuron an ORF (Figure 2b, see Methods). The parameters of the ORF simulations were chosen to qualitatively reproduce the off-target effect: stimulating at proximal locations outside of the soma was often sufficient to drive activity.

**Figure 2:**
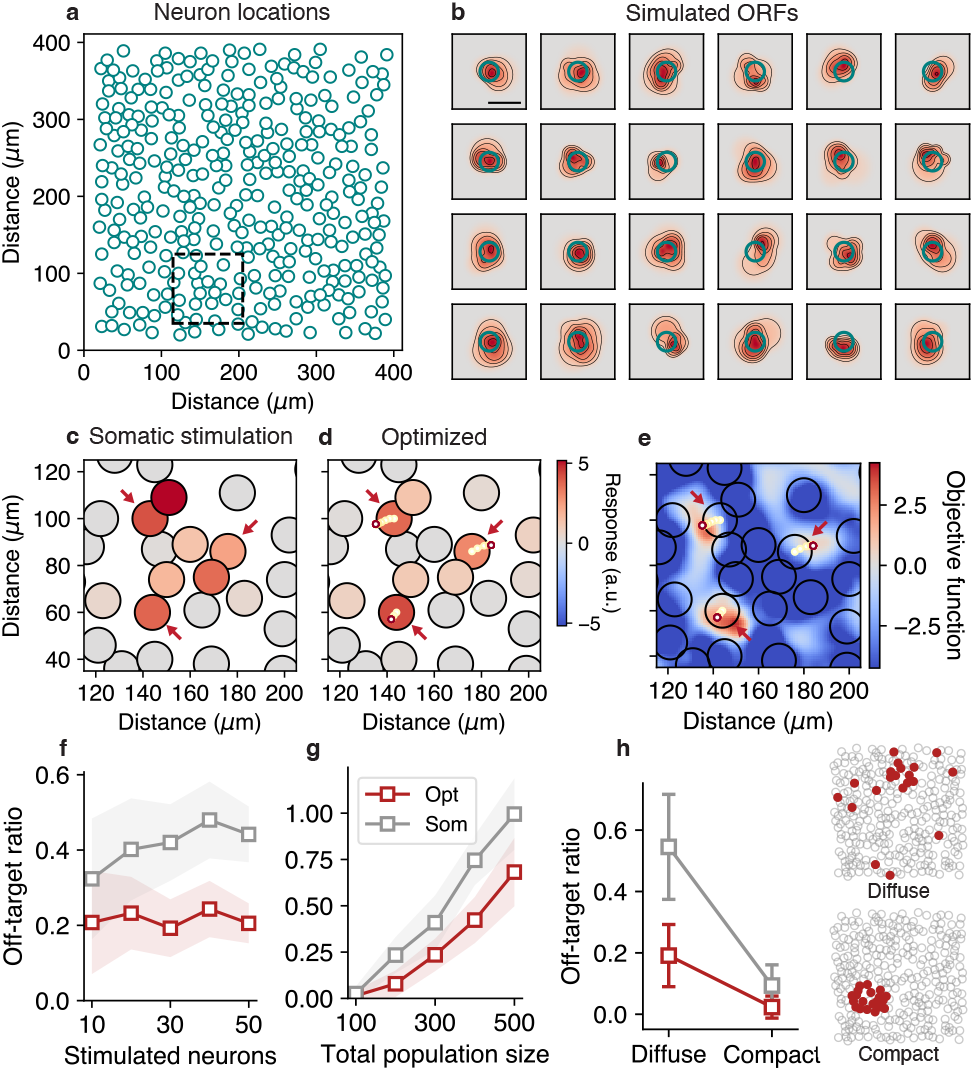
Evaluating NBFR performance using simulations. **a**, We performed simulations involving hundreds of neurons with heterogeneous ORFs to determine the behavior of NBFR. **b**, Simulated ORF parameters were chosen to yield stimulation responses even when targeting proximal locations outside of the soma, qualitatively recreating *in vivo* neuron responses. Scale bar, 20 *µ*m. **c, d**, Comparison of conventional (i.e. “somatic”) stimulation (**c**) with optimized stimulation (**d**). Large circles with black edges represent neurons, small circles with colored edges show optimization path. In this example, the three neurons indicated by the red arrows are stimulated simultaneously. Unlike optimized stimulation, somatic stimulation leads to substantial off-target activation. **e**, Objective function surface shows how optimization repositions holographic stimulation sites based on collective ORF information from the local population of neurons. Note that color corresponds to the objective function based on ground truth ORFs, whereas optimized stimulation sites are based on the learned ORFs. Hence optimized targets (e.g. bottom left stimulus) may not necessarily align with the true global optimum due to constraints on ORF learning from limited or noisy data. **f-h**, Performance of NBFR-optimized stimulation (red) compared to somatic stimulation (gray), as a function of the stimulated ensemble size (**f**), the total size of the population (reflecting density of opsin-expressing neurons) (**g**), and whether stimulated neurons are diffuse or compact (**h**). Ensemble size used for **g** and **h**, 20. Error bars represent mean *±*1 standard deviation over 50 simulations.

When performing conventional stimulation by positioning holograms over the centroids of the target neurons’ somas (“somatic stimulation”), the spatial extent of the ORFs caused frequent off-target activation of multiple non-target neurons in the immediate proximity of the targets (Figure 2c). However, first mapping the ORFs using NBFR enabled the repositioning of the stimulation sites so as to suppress off-target effects (Figure 2d). The optimization algorithm seeks to maximize an objective function given by the sum of the evoked activity in the target neurons, minus the evoked activity in the non-target neurons (Methods). Thus, to understand why these specific sites were chosen, we visualized the objective function surface and overlaid the neuron locations, which revealed how the optimization traced a path of stimulation sites that integrated ORF information from all local neurons (Figure 2e) to avoid off-target activation.

Next, we evaluated the performance of NBFR while systematically varying key variables over a range of values that resembled typical experimental conditions (Figure 2f-h). To quantify stimulation accuracy, we used a value we call the “off-target ratio”: the number of activated non-target neurons per activated target neuron. In our simulations, we found that the number of simultaneously stimulated neurons did not markedly affect the off-target ratio (Figure 2f), with NBFR-based stimulus optimization having ~50% fewer off-targets per target when stimulating 10 to 50 neurons at once (off-target ratio for somatic stimulation, ~0.4; optimized, ~0.2). However, whether targeted ensembles were composed of neurons that were spatially compact (“compact ensembles”) or diffuse (“diffuse ensembles”) did affect the off-target ratio. While stimulating compact ensembles (number of constituent neurons, 20) did not create a sizable difference in off-target ratio between somatic vs optimized stimulation, stimulating diffuse ensembles with NBFR-optimization resulted in half the number of non-targets per target compared to somatic stimulation (Figure 2h; ~4 off-target neurons with optimization vs ~9 with somatic stimulation for a 20-target ensemble). Hence, optimization achieves maximal utility when stimulating diffusely spaced ensembles of neurons, since off-target activation in compact ensembles is likely to primarily affect other nearby members of the target ensemble.

We also found that NBFR performance depends on the cell-packing density. To assess this, we fixed the size of the stimulation FOV while varying the total size of the population. As expected, for sparse populations of neurons (e.g. 100 neurons in a 411×411 *µ*m^2^ FOV) off-target activation was rare. However, as the number of neurons increased, the performance gap between somatic and optimized stimulation widened (Figure 2g), demonstrating the increasing need for stimulus optimization tools in experiments with high opsin-expression density.

### Photostimulus optimization can be performed in real time using NBFR

We designed our approach to both support real-time experiments and to be highly modular. As such, we implemented NBFR in Python using only standard scientific computing libraries such as NumPy, and made use of highly efficient non-negative least squares solvers (Methods). To further reduce runtime requirements when mapping ORFs, our proposed experimental design is to use single-trial responses to stimulation instead of averaging across repetitions. Then, in order to gain statistical power, we make use of the fact that responses to stimulation tend to be correlated in space due to the spatial structure of ORFs. Thus, NBFR will “average away” the uncorrelated noise across nearby stimulation sites while preserving spatially correlated signals that arise from the ORF. Additionally, stimulation sites allocated to (e.g.) neuron *j* during the ORF mapping phase can also provide information about neuron *i*’s ORF if neurons *i* and *j* are close enough. However, if it proves necessary in practice, additional statistical power could be gained by performing multiple repetitions of each stimulus at the cost of a longer ORF mapping phase (see Figure 4e below).

To evaluate suitability for real-time experiments, we next performed benchmarking simulations (Figure 3a). First, we determined the number of stimulation patterns required to probe ORFs from increasingly larger populations. For ORF mapping using 10-target stimulation performed at 1 Hz, our experimental design typically required fewer stimulation trials than the number of neurons being mapped – a sublinear scaling behavior similar to compressed sensing. This corresponded to an experiment time of less than just 4 minutes to probe 500 neurons (Figure 3b). Next, we characterized the time required to fit NBFR models to the neural population responses. We found that model fitting was extremely fast: ranging from just 1-3 milliseconds per neuron on average for populations of 100 to 500 neurons (total model fitting time, 0.1 to 1.4 seconds; Figure 3c). Finally, the time required to optimize stimulation patterns was an increasing function of both the total population size and the size of the stimulated ensemble, but did not exceed 600 milliseconds on a Macbook Pro (2023 Apple M3 Pro, 36 GB RAM) under typical experimental conditions (Figure 3d,e). Together, these results establish NBFR’s suitability for real-time experiments.

**Figure 3:**
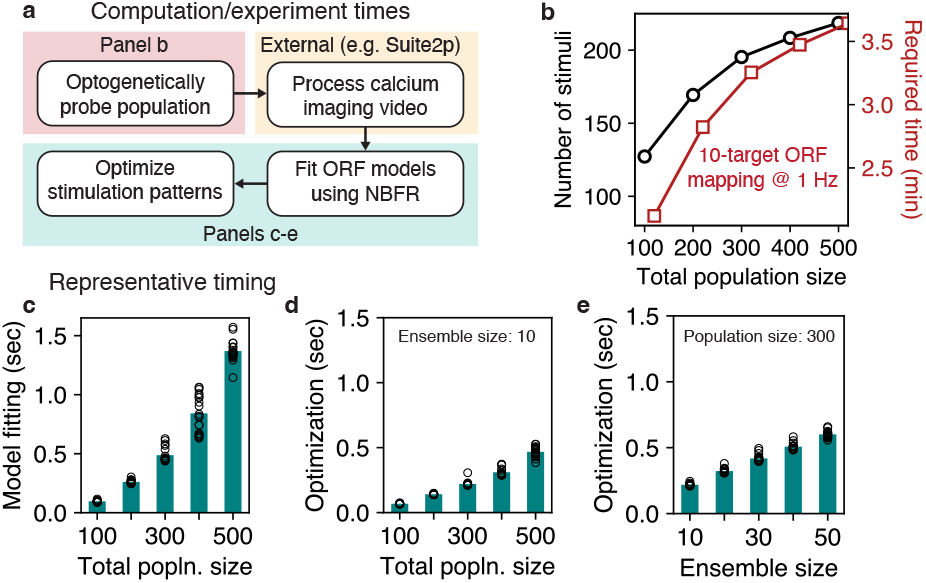
Time required to use NBFR in practice. **a**, Overview of steps involved in NBFR-based photostimulus optimization. The processing of calcium imaging videos is performed by external software such as suite2p [Pachitariu et al., 2016], PMD [Buchanan et al., 2018], or CaImAn [Giovannucci et al., 2019]. **b**, ORF mapping is performed in parallel, typically requiring fewer stimuli than neurons (black line). Further, ORF mapping can be performed in just a few minutes assuming stimulation is performed at 1 Hz (red line). **c**, Time required to fit NBFR models to all neurons. Each data point corresponds to a different simulated FOV. **d-e** Time required to optimize stimulation patterns given ORF models from **c**, as a function of number of total population size (**d**) and stimulated ensemble size (**e**). Each data point in panels **d** and **e** corresponds to a different randomly chosen stimulation pattern (one pattern per simulated FOV).

### Photostimulus optimization is robust to local recurrent excitation under typical conditions

In some cases, two-photon optogenetic stimulation might generate activity in neurons that are untargeted (in terms of both direct and off-target optical excitation), but that are synaptically connected to one or more target neurons. Ensemble stimulation could therefore engage recurrent interactions that have the potential to contaminate ORF mapping, and consequently compromise stimulus optimization. However, in many brain regions excitatory connectivity among pyramidal neurons is extremely sparse [Holmgren et al., 2003], and in hippocampal area CA1 (where our *in vivo* experiments take place, see below) has been reported to be *<*1% [Knowles and Schwartzkroin, 1981, Thomson et al., 1996]. Instead, ensemble stimulation in CA1 appears to strongly engage local inhibition rather than monosynaptic excitation [Robinson et al., 2020, Rolotti et al., 2022], and any polysynaptic excitation that might occur would likely be weak compared to the large, direct photocurrents evoked by two-photon optogenetic stimulation [Sridharan et al., 2022].

Nevertheless, we sought to characterize how such recurrent interactions could impact stimulus optimization (Figure 4a). We hypothesized that ORF mapping could be robust to this source of variability because occasional recurrent excitation could simply be “folded into” the noise component in the NBFR model unless the recurrent effects were unusually strong.

**Figure 4:**
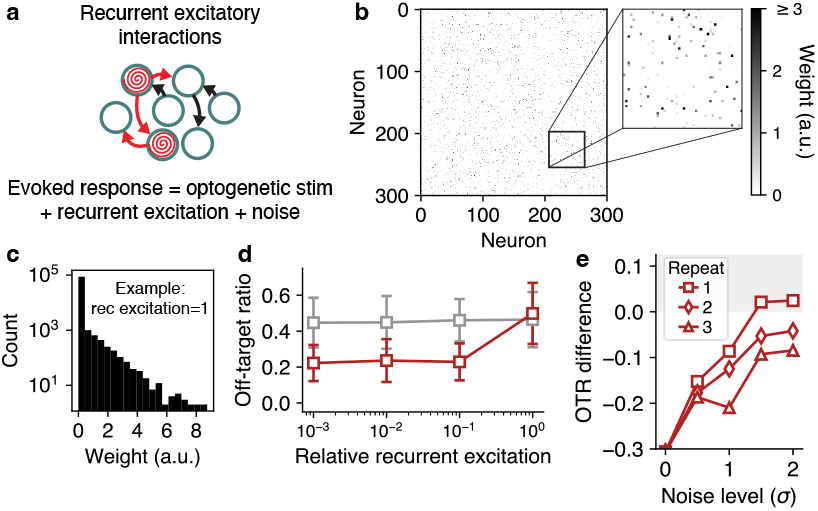
Impact of recurrent excitation and imaging noise on stimulus optimization. **a**, Recurrent excitatory connections could be engaged by ensemble stimulation during ORF mapping. **b**, Example simulated pyramidal-pyramidal functional connectivity matrix with a connection probability of 0.05 and average connection strength of 1. **c**, Heavy-tailed (exponential) distribution of connection strengths corresponding to connectivity matrix in panel **b. d**, Off-target ratio as a function of increasing recurrent excitation levels during ORF mapping. Recurrent input disabled when evaluating off-target effects to isolate impact of direct OTS. Recurrent excitation only compromises evoked neural activity when connection strengths are approximately as large as the optogenetic stimulation (ORFs normalized such that their amplitude is 1). Error bars show one standard deviation over 50 ensembles. **e**, Off-target ratio of optimized stimulation relative to somatic stimulation (i.e. somatic *−* optimized), as a function of increasing standard deviation of the imaging noise (*σ*). Shaded gray region indicates where somatic stimulation is equivalent to, or better than, optimized stimulation. Under high noise levels, performing multiple repetitions of the ORF mapping protocol can preserve the utility of NBFR. Each data point in **d** and **e** is an average over 50 test ensembles, where each ensemble is sampled from a new, randomly initialized simulation.

We performed simulations (see Methods) with a relatively high “functional” pyramidal-pyramidal connection density of 5%, under the assumption that such functional connections represent polysynaptic pathways mediated by unobserved interneurons. By systematically varying the distribution of connection strengths in our simulations (Figure 4b,c), we found that, indeed, recurrent excitation only constrained stimulus optimization when the average functional connection strength approached the magnitude of the direct photostimulation itself (Figure 4d), at which point the off-target ratio resulting from optimized stimulation was equal to that of somatic stimulation. However, this is extremely unlikely to occur in practice as monosynaptic connection strengths in hippocampus and cortex are typically orders of magnitude smaller than photocurrents elicited by direct optogenetic stimulation [Thomson et al., 1996, Antin et al., 2024, Triplett et al., 2025]. Thus, our results suggest that recurrent excitation should not meaningfully disrupt stimulus optimization under standard experimental conditions.

### *In vivo* experiments demonstrate the utility of NBFR

To demonstrate our method in a real-world setting, we applied it to holographic stimulation experiments in hippocampal area CA1 in awake, behaving mice (Figure 5a). We used a bicistronic virus to simultaneously express the calcium indicator GCaMP6m and excitatory opsin ChRmine in CA1 pyramidal neurons. This approach guaranteed co-expression of the opsin and calcium indicator in transduced neurons and therefore allowed us to accurately estimate off-target effects in all neurons responsive to stimulation. Next, to excite target neurons, we used a spatial light modulator to form a pattern of diffraction-limited spots and “spiral scanned” them over the target neurons’ somas simultaneously (Figure 5b,c), a standard and widely-used approach for holographic stimulation experiments [Rickgauer and Tank, 2009, Emiliani et al., 2022].

**Figure 5:**
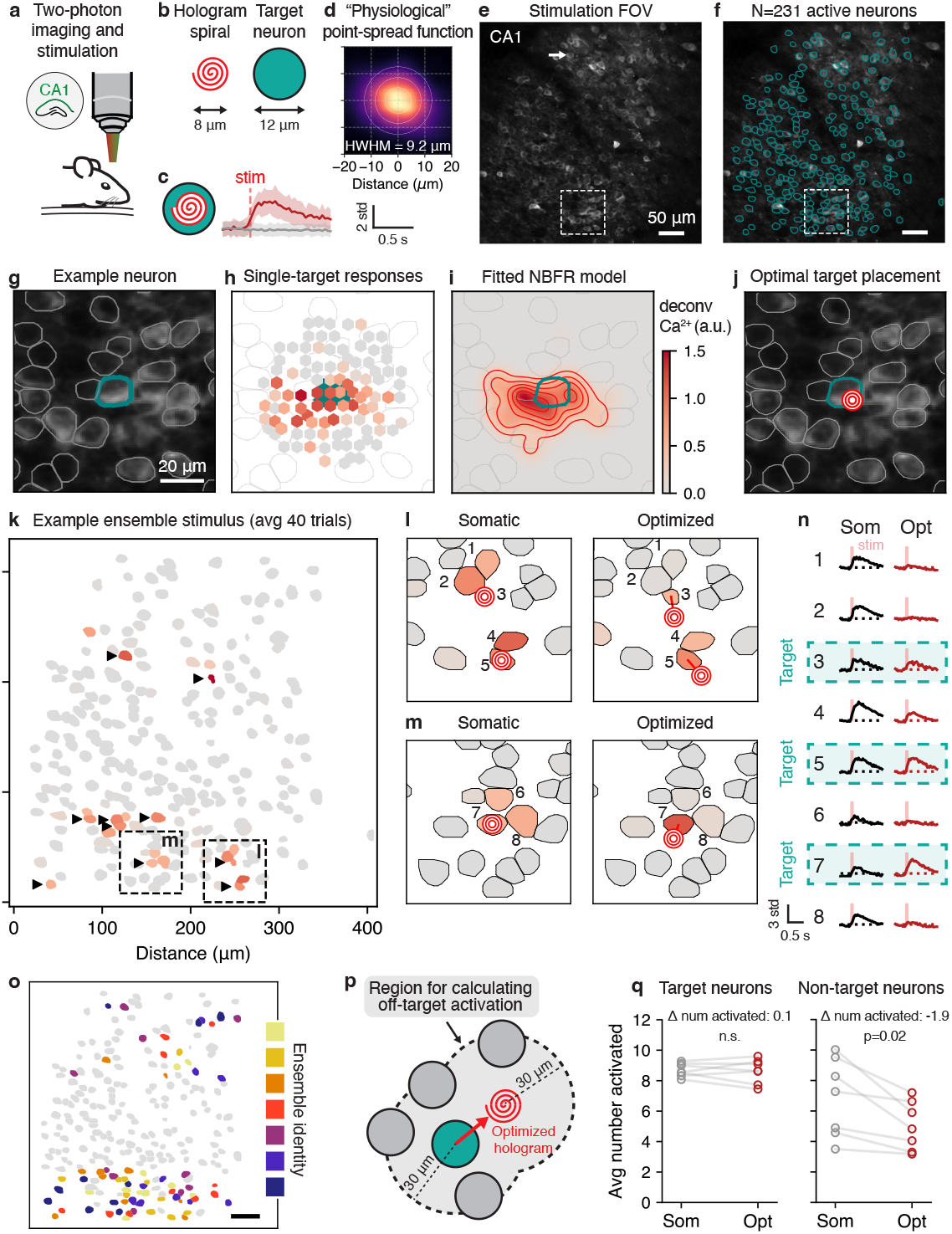
Real-time optimization of holographic stimulation sites can mitigate off-target stimulation *in vivo*. **a-c**, Experimental setup for two-photon holographic stimulation and calcium imaging in CA1. **d**, Physiological point-spread function estimated by averaging all neurons’ responses at proximal stimulation sites. **e**, CA1 stimulation FOV for an example experiment. White arrow corresponds to example neuron in **g. f**, N=231 neurons were segmented using Suite2p. **g-j**, Location, responses to single-target stimulation, inferred ORF, and optimized stimulus for an example neuron. **k**, Example holographic ensemble stimulation pattern. Target neurons (indicated by black triangles) stimulated at their somas generate off-target activity. **l, m**, Comparison between somatic and optimized stimulation shows that off-target activation can be avoided while still activating target neuron. Cf. dashed windows in panel **k. n**, dF/F traces for somatic and optimized stimulation for the 8 example neurons shown in panels **l** and **m** confirms reduction in off-target stimulation. **o**, Locations of all stimulated ensembles in this experiment. Color indicates ensemble identity. Scale bar, 50 *µ*m. **p**, To calculate off-target effects, the activity of all neurons within 30 *µ*m of either the target neuron or the optimized stimulus are considered. **q**, Effect of stimulus optimization for the 7 ensembles from panel **o**. Activated neuron determined by thresholding the z-scored calcium response for all target or non-target neurons. While the activity of the target neurons did not significantly change with optimization, the number of activated non-target neurons reduced by 2 (p=0.02, dependent *t*-test). All stimulation patterns were repeated 40 times each.

We then performed an ORF mapping phase by defining grids of stimulation sites surrounding each neuron, and targeted randomized sets of ten such sites for stimulation at a time (similar to our simulations, cf. Figure 1a; Supplementary Figure 1) while the animal (head-fixed) was free to walk on a wheel. After collecting the population responses to ensemble stimulation, we examined the neural activity of each neuron when only one out of ten holographic stimulation sites was within 30 *µ*m of the target neuron, thus approximating the mapping of ORFs that would be obtained using single-target stimulation only (Figure 5g,h). Averaging these pseudo single-target responses from every neuron in the FOV yielded an average “physiological” point spread function (PPSF) with a half-width at half-max of 9.2 *µ*m (Figure 5d-f), consistent with previous reports on the resolution of “all-optical” systems [Pégard et al., 2017, Shemesh et al., 2017, Triplett et al., 2025].

However, individual neurons could often be stimulated surprisingly far outside of the average PPSF; in some cases 15 *µ*m or more from the cell centroid in the radial axis (e.g. Figure 5h; also see [Triplett et al., 2023] for similar results obtained using loose-patch recordings), despite the hologram spiral having a diameter of just 8 *µ*m (Figure 5b). Thus, we found that individual variability in ORFs is masked by averaging when obtaining conventional PPSFs.

We used NBFR to infer the ORFs of all 231 neurons responding to photostimulation in the FOV (see Supplementary Figure 2 for examples). Obtaining these ORFs enabled the computation of optimal stimulation sites for target neurons while accounting for the ORFs of neighboring non-target neurons (Figure 5j).

Next, we applied our method to optimize the targeted activation of neural ensembles. We selected ensembles for testing based either on the quality of their inferred ORFs or by the expected improvement in precision resulting from optimization (Methods). Conventional somatic stimulation resulted in target neurons being activated with high probability, but also recruited neighboring non-target neurons due to overlapping ORFs (Figure 5k). Optimizing the exact holographic target placements based on NBFR-inferred ORFs resulted in substantial reduction of off-target activation (Figure 5l-n), and in some cases even increased the activation of the target neurons despite repositioning of the spiral away from the target neuron centroid (Figure 5n, “Somatic” vs “Optimized”). Over 7 ensembles within the same FOV (Figure 5o, Supplementary Figure 3), target optimization resulted in a reduction of ~2 non-target neurons per 10-target ensemble stimulus (change in average number of activated non-target neurons over 40 repetitions, −1.9; p=0.02, dependent *t*-test), with no statistically detectable difference in the activation of the target ensembles (change in average number of activated target neurons, 0.1; n.s., dependent *t*-test; Figure 5p,q). Thus, NBFR-based target optimization can reduce off-target activation under *in vivo* conditions.

### Motion is the primary limiting factor for NBFR-based stimulus optimization *in vivo*

To characterize the performance of *in vivo* stimulus optimization, we repeated the experiment for a total of 47 ensembles across 7 FOVs (from 4 mice; Figure 6a, Supplementary Figure 4). On average, optimization resulted in 3 fewer non-targets activated per 10-target ensemble (change in average number of target neurons activated, −3.1; p<10^−9^, dependent *t*-test) with an average reduction in target neuron activity of just 1 per ensemble (change in target neurons activated, −1.1; p<10^−9^, dependent *t*-test).

**Figure 6:**
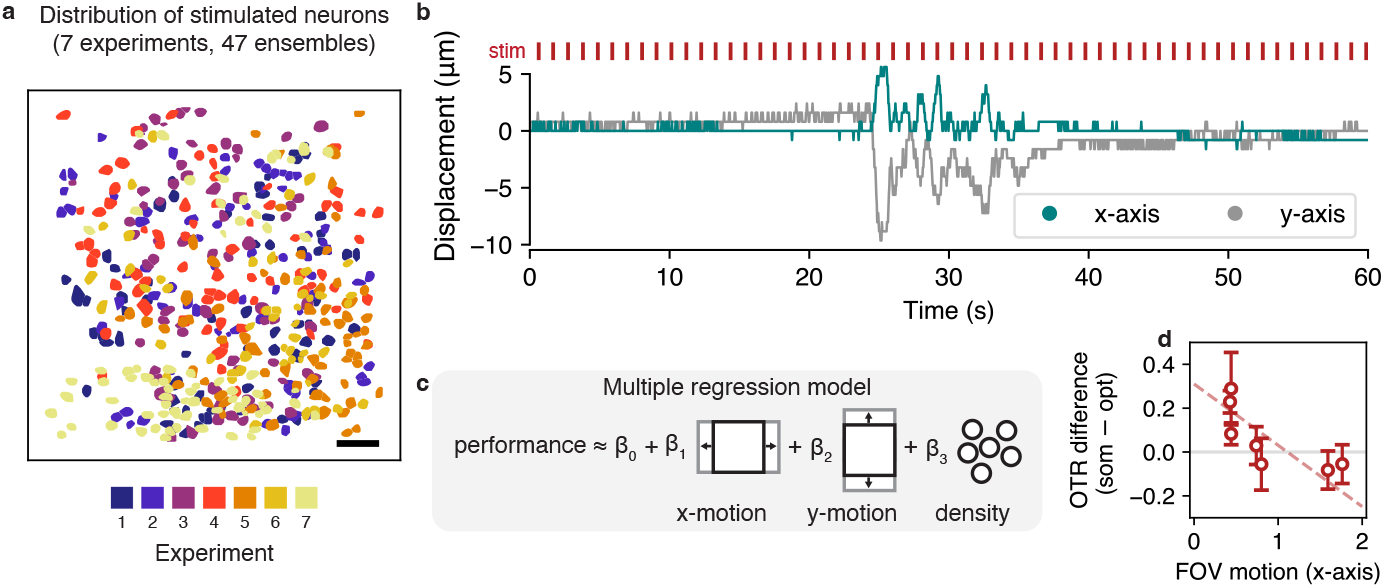
Motion constrains performance improvement of NBFR-based stimulus optimization. **a**, Stimulated ensembles across all 7 experiments (*n*=47 ensembles, 4 mice). Ensembles colored by experiment. Scale bar, 50 *µ*m. **b**, Example FOV motion correction traces from NoRMCorre. **c**, Multiple linear regression model used to determine experimental parameters most predictive of algorithm performance (parameters considered, x-motion, y-motion, and nearby cell density). **d**, Multiple regression identifies FOV motion in the x-axis (defined as the average absolute value of the motion correction trace) as the primary constraint on the performance improvement of NBFR-based optimization *in vivo*. Each data point corresponds to one experiment, error bars show standard error over 5-7 test ensembles per experiment.

We hypothesized that the two factors most likely to affect algorithm performance would be (1) the prevalence of nearby opsin-expressing neurons, and (2) FOV motion resulting from movement of the animal (despite a head-fixed preparation, Figure 6b). We therefore calculated the difference in the off-target ratio between optimized and somatic stimulation trials (the “OTR difference”), and performed a multivariate regression of the OTR difference onto both the number of nearby non-target neurons and FOV motion (Figure 6c). While the number of nearby non-target neurons was not a statistically significant predictor of OTR difference (p=0.06), the motion energy coefficient – particularly in the x-axis – was significant (motion in x-axis, p=0.01; y-axis, p=0.27). In experiments where motion energy was low (~0.4 *µ*m FOV displacement per frame on average), the OTR difference was the largest (difference between off-target ratio for optimized stimulation and somatic stimulation, 0.2 off-targets per target; Figure 6d). However, the OTR difference was minimal for experiments with high motion energy, indicating that the utility of target optimization is primarily constrained by FOV motion.

## Discussion

Overcoming OTS is critical for precise causal perturbation experiments. We have introduced a new approach based on computational optimization that enables a precision potentially higher than what could be achieved by combining two-photon excitation and soma-targeted opsins alone. Importantly, this improvement does not require changes to the optical hardware or opsins, making NBFR a practical addition to existing holographic systems. We expect that our approach could represent an important tool for an array of critical neuroscience experiments. For example, OTS creates ambiguity about the presynaptic origin of a postsynaptic current in synaptic connectivity mapping [Packer et al., 2012, Triplett et al., 2025], prevents the optogenetic recreation of observed neural activity patterns in “playback” experiments [Grosenick et al., 2015], and can compromise stimulation experiments where neurons with distinct functional roles are densely intermingled but only one type must be activated (e.g., stimulation of cortical neurons tuned to a specific visual orientation when orientation selectivity follows a “salt-and-pepper” organization [Kaschube, 2014], or stimulating place cells in CA1 [Robinson et al., 2020]). Our real-time optimization approach could be an essential tool for reducing the prevalence of these effects, potentially enabling experiments that would otherwise be precluded by OTS.

From a technical standpoint, the ability to fit hundreds of ORF models in just a few seconds on a laptop is a major advance over previous work [Triplett et al., 2023], which relied on intensive matrix inversions to implement Gaussian process regression. Here, once ORFs are estimated, optimizing holograms requires just a few hundred milliseconds using standard Python libraries, highlighting the practical feasibility of integrating NBFR into closed-loop experimental pipelines.

Despite these advances, we encountered some limitations with NBFR-based stimulus optimization. In particular, when the risk of OTS is particularly high (e.g., due to many highly responsive non-target neurons surrounding a target neuron), optimization may recommend aborting stimulation altogether in order to maximize its objective function (i.e. by repositioning the stimulation site to an area where there are no opsin-expressing cells). Although this is a rational outcome of the optimization, it could be problematic in experiments where target activation should be guaranteed. Future work could explore integrating neuron-specific constraints into the optimization algorithm to enforce a minimum level of target activation while still avoiding OTS.

Further, while fast, our current method only optimizes hologram coordinates in 2D. Extending this framework to jointly optimize 3D coordinates and per-cell laser power (as with Bayesian target optimization) would be an important next step to achieving the highest possible accuracy [Triplett et al., 2023]. However, doing so will require a new optimization approach that balances the flexibility of Bayesian target optimization with the scalability and efficiency of NBFR. In addition, it would also require sufficient control over the stimulation system; specifically, independent control over the 3D positioning and laser power of multiple holograms simultaneously, which not all commercial holographic microscopes currently support. One potentially useful intermediate step would be to combine 2D stimulus optimization with 3D imaging – a configuration that may be more accessible than joint 3D stimulation and 3D imaging. This would allow for monitoring of off-target stimulation in the axial dimension and could be straightforwardly achieved by extending NBFR’s basis functions to 3D while leaving the optimization routine as-is.

Another important direction will be to adapt NBFR-based stimulus optimization to support voltage imaging experiments [Hochbaum et al., 2014], which offer exceptional temporal resolution but also face numerous technical challenges when combined with photostimulation, especially in the two-photon regime. In principle, the ORF mapping phase could be completed extremely quickly due to not needing to wait for the comparatively slower calcium indicator kinetics. However, cross-talk between stimulation and imaging in two-photon mode currently precludes simultaneous expression of the opsin and sensor in the same neurons. One-photon experiments do not face this limitation to the same extent, but have significantly degraded spatial resolution that computational approaches may not be able to overcome. Precise photostimulus optimization for joint two-photon stimulation and voltage imaging experiments will therefore likely require the engineering or discovery of new opsin-sensor combinations that are sufficiently spectrally orthogonal in the two-photon regime.

Additionally, nonstationarities are pervasive *in vivo*, and it is possible that ORFs could change over time (e.g. with behavioral state). While our model assumes a static opsin-expression profile over the soma and proximal dendrites, overall changes in intrinsic excitability could require multiplicative adjustments to the ORF amplitude [Triplett and Goodhill, 2022]. We hypothesize that this could be straightforwardly done “online” by scaling the ORF model to match the amplitude of recently evoked calcium transients.

More broadly, maintaining precision as all-optical tools continue to scale to larger populations will require adapting stimulation patterns automatically based on learned response properties, as manual adjustment will be increasingly impractical. Thus, we expect that approaches like NBFR will be a critical part of a growing toolbox of computational methods [Eybposh et al., 2020, Draelos and Pearson, 2020, Triplett et al., 2023, Navarro and Oweiss, 2023, Antin et al., 2024, Wagenmaker et al., 2024, Triplett et al., 2025, Draelos et al., 2025] aimed at optimizing holographic stimulation experiments for neural circuit analysis.

## Acknowledgements

MAT and EB thank the organizers of the Cajal Advanced Experimental Neuroscience Training Programme at the Champalimaud Centre, where this collaboration was initiated. MAT is supported by NIH award K99NS135649. EB was supported by a fellowship from Boehringer Ingelheim Fonds. MAT and LP are supported by the Kavli Foundation and the Gatsby Charitable Foundation (GAT3708). LP is supported by DoD OUSD (R&E) under Cooperative Agreement PHY-2229929 (The NSF AI Institute for Artificial and Natural Intelligence). AP is supported by the Engineering and Physical Sciences Research Council (awards EP/R513143/1 and EP/W524335/1). DSP is supported by NIH award R01EY033950 and Zuckerman Institute Team Science Funds. MH is supported by grants from the Wellcome Trust (PRF 201225 and 224688), ERC (AdG 695709), MRC (MR/T022922/1) and the BBSRC (BB/N009835/1).

## Code availability

Code for this manuscript is available at https://github.com/marcustriplett/photostim-optimizer.

## Methods

### Animals

All experimental procedures were carried out under license from the UK Home Office in accordance with the UK Animals (Scientific Procedures) Act (1986). All experiments used adult C57BL/6 WT mice (Charles River Laboratories) of both sexes aged between 6-18 weeks before the first surgery. Animals were single-housed in an enriched environment and kept on a 12-hour light/dark reverse cycle.

### Surgical procedures

All surgical procedures were performed under isoflurane anaesthesia (2%, Isoflo) with additional analgesia (buprenorphine – Vetergesic, meloxicam – Metacam) using an automated stereotaxic system. Following surgery, animals were orally administered meloxicam (Metacam) for three days post-operatively. Animals underwent two consecutive surgeries. We performed an initial viral injection surgery using a bicistronic virus (AAV8-CaMKIIa-GCaMP6m-p2A-ChRmine-Kv2.1-WPRE virus (2.75 × 1012 *µ*g/mL; Stanford Neuroscience Gene Vector and Virus Core, GVVC-AAV-180, [Marshel et al., 2019])) in dorsal CA1 neurons. The virus was first diluted to a range of 1:2 – 1:10 from stock using buffer solution (20 mM Tris,140 mM NaCl, 0.001% Pluronic F-68, pH 8.0). Next, a small (0.5 mm diameter) craniotomy was made above the right hippocampus at −1.85 mm AP, 1.45 mm ML from bregma. A glass-pipette (Drummond Scientific Company, Wiretrol II, 5-000-2005) was connected to a hydraulic injection system (Narishige MO-1) loaded with diluted virus and carefully lowered to −1.4 mm DV from bregma. A total volume of 1 *µ*l was slowly injected at 100 nl/min.

Following a 1-week recovery period, a second surgery was performed to obtain optical access in CA1 as described previously [Robinson et al., 2020]. In brief, following a craniotomy centered above the initial injection site, cortical tissue above the corpus callosum was carefully aspirated under continuous irrigation with cold aCSF (4^*°*^ C). Next, a glass window attached to the bottom of a custom-made metal cannula (1.8 mm height, 3 mm diameter) was carefully lowered into the craniotomy. The gap between the cannula and skull was sealed using tissue adhesive (Dermafuse). Finally, a custom-made titanium head plate was affixed to the skull using dental cement.

### Two-photon calcium imaging with simultaneous holographic optogenetic stimulation

Simultaneous 2P holographic stimulation and imaging were performed as described previously [Robinson et al., 2020] using a Bergamo II rotating system (Thorlabs) equipped with a spatial light modulator (Thorlabs) and 10x air-immersion objective (0.5 NA, 7.8 mm WD, Thorlabs). Experiments were controlled using ThorImage (v4.2, Thorlabs). For calcium imaging, 930 nm light delivered by a Ti:sapphire laser (Chameleon Ultra II, Coherent) was focused onto the CA1 pyramidal cell layer at a 5 degree angle. An average imaging power of 30 mW was used across a 600×600 *µ*m FOV at 512×512 pixel resolution, resulting in an average frame rate of 30 Hz. Emitted fluorescence was collected through a 562 nm long pass dichroic and a 525-550 nm bandpass filter, after which it was amplified by photomultiplier tubes (Hamamatsu). Simultaneous 2P holographic optogenetics was achieved through a secondary light path. Holograms were calculated using a Gerchberg-Saxton algorithm and applied to a reflective multilevel spatial light modulator (OverDrive Plus, Meadowlark, 7.68×7.68 mm active zone at 512×512 pixels) used to shape a 1 MHz rep rate 1030 nm femtosecond fiber laser (BlueCut 10, Menlo Systems) into multiple target beamlets. Imaging and stimulation beams were aligned using a 740 nm short pass and a 1050/40 nm bandpass dichroic before reaching the back aperture of the objective. The resulting patterns were spiralled over an 8 *µ*m area centred on the targeted neurons using a galvo-galvo scanning system at 5 mW per cell. For ensemble activation, neurons were targeted with 5 consecutive 20 ms long spirals for 100 ms total stimulation time. Stimulus identities were randomly interleaved. Before each experiment, a software calibration step was performed by burning spots on a fluorescent slide and calculating the required affine transformation between the SLM pattern and output at the imaging plane to ensure maximum targeting accuracy.

### Experimental design for ORF mapping and stimulus optimization

To map ORFs, we first extracted the coordinates of all neurons in the imaging FOV using suite2p [Pachitariu et al., 2016]. Next, we positioned a grid of stimulation sites over the centroid of each neuron, covering a −16 to 16 *µ*m range (relative to the cell centroid) with a step-size of 8 *µ*m, yielding 25 stimulation sites per neuron. We then removed stimulation sites that were within 5 *µ*m of any other stimulation sites associated with other neurons’ grids (removal performed sequentially in a randomized order). When the density of opsin-expressing neurons in the FOV is low, the number of stimulation sites is high relative to the population size. However, as density increases, the stimulation grids increasingly overlap. Thus, when nearby redundant points are removed, the number of stimulation sites becomes progressively smaller relative to the population size, and ORF mapping becomes highly efficient (cf. Figure 3b).

Subsequently, we restructured the set of stimulation sites into trials of 10-target ensemble stimulation patterns. Because we stimulated at a high speed relative to the calcium indicator decay time, we aimed to avoid returning to stimulate in the proximity of any given cell too quickly; otherwise, the evoked calcium transients would not have time to return to baseline and our ORF estimates would be artificially elevated. To achieve this, we randomly selected an ensemble stimulus for the first trial. Then, for any subsequent trial, we randomly selected stimuli until each constituent target was a minimum distance away from every target on the previous trial. The resulting sequence of stimulation patterns was then used to probe the neurons’ ORFs. GCaMP-expressing neurons that did not respond to stimulation during our mapping phase were deemed not excitable and excluded from analysis.

After the ORFs were mapped, neurons selected to participate in test ensembles were chosen either manually by inspecting the quality of their inferred ORFs (2/7 experiments), or as those estimated to show the largest expected improvement from stimulus optimization (estimated by performing the optimization and sorting neurons by largest improvement in objective function, 5/7 experiments). No major difference in algorithm performance between ensemble selection methods was identifiable during post hoc analysis. Once ensembles were selected, the gradient descent step-size and the number of gradient descent iterations were tuned in real time based on whether the objective function was determined to have converged. The resulting stimulation sites were then used to evaluate the performance of stimulus optimization relative to somatic stimulation.

### Adaptive non-negative basis function regression

Here we describe NBFR – the computational framework used to estimate ORFs. We model the response *y*_*nk*_ ∈ ℝ to an optogenetic stimulus **x**_*k*_ ∈ ℝ^*J×*2^ on trial *k* = 1, …, *K* as the summation of activity evoked by *J* different holographic target spirals (though the approach is also suitable for scanless illumination techniques [Papagiakoumou et al., 2020, Pégard et al., 2017]),

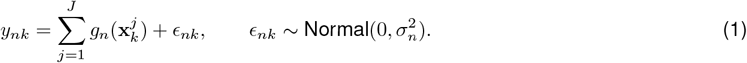

Here *g*_*n*_: ℝ^2^ → ℝ is the ORF, which specifies the calcium-dependent fluorescence signal evoked by stimulating at any given coordinate near the neuron. The key idea is to model the ORFs using non-negative basis function regression (NBFR). To this end, for each neuron *n* we define a collection of basis functions 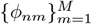 defined as

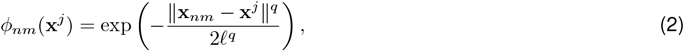

where each **x**_*nm*_ ∈ ℝ^2^ (for *m* = 1, …, *M*) is the location of a stimulus within some distance *d* (typically 30 *µ*m) from the centroid of neuron *n* and where ℓ is the length scale of the basis function. This model is “adaptive” in the sense that the set of basis functions is determined by the stimulation sites: the model places down a basis function at any location that was stimulated by any member of an ensemble stimulus that was sufficiently nearby, and thus the basis set varies from experiment to experiment depending on the spatial organization of neurons. The exponent *q* ≥ 2 determines how sharp the basis function is: if *q* = 2 then *ϕ*_*nm*_ corresponds to the standard radial basis function, and if *q >* 2 then *ϕ*_*nm*_ is known as a “super-Gaussian” function, with a flatter top and sharper drop-off.

We then define the ORF *g*_*n*_ as a weighted sum of such basis functions,

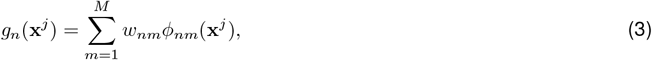

where each *w*_*nm*_ ∈ ℝ_*≥*0_ is non-negative. As it turns out, non-negativity of the basis function weights is critical; otherwise, responses to novel (i.e., out of sample) holograms are frequently predicted as implausibly negative.

We use data from an initial calibration session to estimate the ORFs. The calibration session yields responses **y**_:,1_, …, **y**_:,*K*_ to holographic ensemble stimuli **x**_1_, …, **x**_*K*_, where each **y**_:,*k*_ ∈ ℝ^*N*^ is the population response to stimulus **x**_*k*_. Given such calibration data, we define a design matrix **Φ**_*n*_ ∈ ℝ^*K×M*^ by specifying each element as

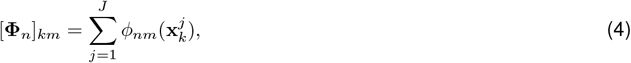

i.e., the contribution of basis function *m* to neuron *n*’s response on trial *k*. Then, NBFR proceeds by solving a non-negative least squares problem to estimate the regression coefficients,

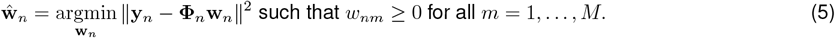

Equation 5 is repeated in turn for each neuron *n*. Note that standard non-negative least squares solvers are extremely efficient – each model requires only several milliseconds to fit for an experiment with several hundred neurons and trials (for details see section *Photostimulus optimization can be performed in real time using NBFR*).

### Stimulus optimization

Given the fitted NBFR model coefficients 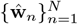, we next turn to optimizing holographic stimulation sites in order to evoke specific neural activity patterns. This requires activating target neurons while avoiding non-target neurons as much as possible. To this end, let Ω ⊆ {1, …, *N*} represent a set of target neuron indices. We propose to optimize the objective function

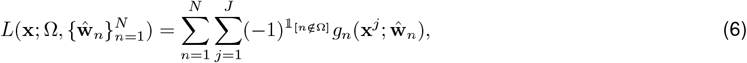

where we have used 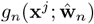 to represent the sum 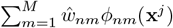. The objective function *L* therefore gives the sum of the evoked activity in the target neurons, minus the evoked activity in the non-target neurons. Thus, we can achieve our desired activity pattern by maximizing *L* as a function of the stimulus parameters **x**.

Note that we can also optimize Equation 6 using an “importance-weighted” gradient descent algorithm. In particular, one can adjust the objective function in Equation 6 such that each neuron *n* receives an “importance scale” *α*_*n*_ ∈ ℝ_*≥*0_ that specifies how important it is to achieve the desired activity level for that neuron. This leads to an importance-weighted objective

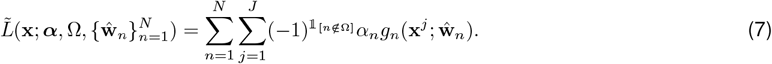

Assuming *q* = 2 in Equation 2 (our typical choice for *q*), we can derive the gradient of Equation 7 with respect to holographic target **x**^*j*^ as

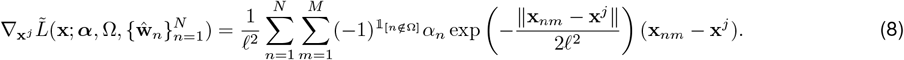

We then perform gradient descent updates using Equation 8.

The presence of the importance weights ***α*** in Equation 8 has some immediate consequences. First, if *α*_*n*_ = 1, the importance-weighted objective 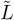 reverts back to the basic objective *L* and all neurons are treated equally. If *α*_*n*_ = 0, then neuron *n* will not be considered at all during the optimization, which may be useful if its ORF appears to have been poorly characterized during the calibration phase of the experiment. Most importantly, by adjusting *α*_*n*_ to take on a sufficiently large value, we can *force* a neuron to be activated or avoided. For example, it may be absolutely essential to activate specific place cells to ensure that an experiment proceeds as required, or to avoid specific place cells in order not to compromise an experiment.

### Simulations

To generate simulations, we first sampled ORFs 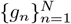 as non-negative mixtures of the basis functions in Equation 2, where the number of constituent basis functions varied, and where each such function had a randomly chosen center and width. We used basis functions with *q* = 4 to form sharper ORFs that qualitatively appeared more consistent with slice recordings [Triplett et al., 2023] compared to a standard radial basis function with *q* = 2.

Next, a simulated neural population response 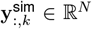 on trial *k* to a stimulus **x**_*k*_ ∈ ℝ^*J×*2^ was obtained by adding noise to the ORF output,

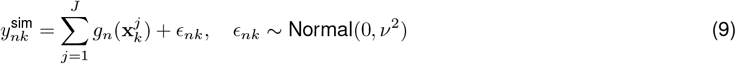

where n = 1, …, N and where ν^2^ is the ground-truth noise variance. NBFR was then used to estimate the ORFs 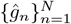 from the noisy simulated data.

To generate neural ensembles that varied in compactness, we used a sampling procedure that weighted the probability of selecting a given neuron based on its distance from a center neuron. Specifically, we first randomly selected a center neuron *n*_c_ from the full population. Then, we selected the remaining members of the ensemble by sampling (without replacement) each neuron *i* with a probability proportional to an exponentially decreasing function of their distance from the center neuron, 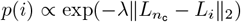. For compact ensembles we set *λ* = 1, and for diffuse ensembles we set *λ* = 0.01.

To evaluate the effect of recurrent functional connectivity, we modelled the neural response to stimulation for neuron *n* as

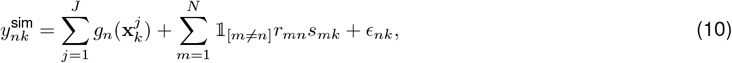

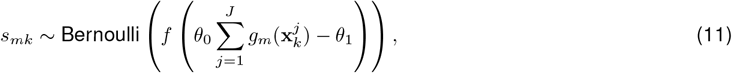

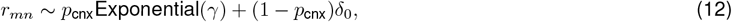

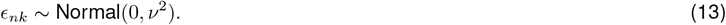

Here *r*_*mn*_ represents the recurrent connection strength from neuron *m* to *n* sampled from a zero-inflated exponential distribution, *s*_*mk*_ represents whether neuron *m* was optogenetically activated and successfully transmitted a spike train on trial *k, f* (*x*) = 1*/*(1 + exp(−*x*)) is the logistic sigmoid function, and *δ*_0_ is the delta function centered at zero. Note that *s*_*mk*_ is a stochastic function of the photostimulation: excitation delivered closer to the ORF peak is more likely to activate the neuron and engage its recurrent pathways.

Default simulation parameters are given in Table 1.

**Table 1.**
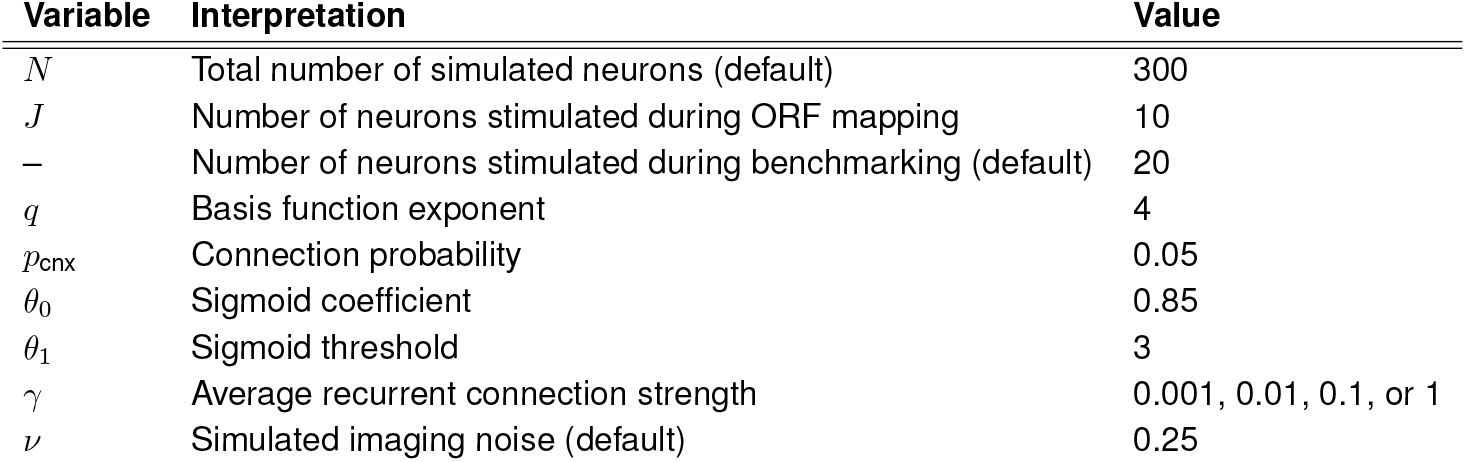
Default simulation parameters.

### Regression model of NBFR performance

To determine the factors underlying the performance improvement of stimulus optimization relative to somatic stimulation we performed a multivariate regression analysis, including motion and local cell density as predictors. The motion energy in the x-axis for a neural recording was defined as

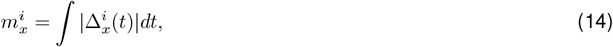

where 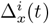 is the motion correction shift applied to calcium imaging video *i* in frame *t*, as estimated using the NoRM-Corre algorithm in suite2p [Pachitariu et al., 2016, Pnevmatikakis and Giovannucci, 2017]. The motion energy 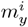 for the y-axis is defined analogously. The “cell density” was simply the number of neurons within 50 *µ*m of any target neuron.Next, we defined a regression model as

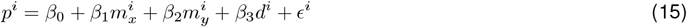

where *ϵ*^*i*^ ~ Normal(0, *ν*^2^) is a Gaussian noise variable and *p*^*i*^ is the performance difference between optimized stimulation and somatic stimulation, evaluated via the off-target ratio

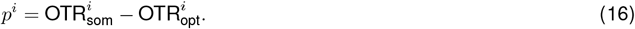

We solved for the regression coefficients using the statsmodels package in Python [Seabold and Perktold, 2010].

## Supplementary Figures

**Figure S1:**
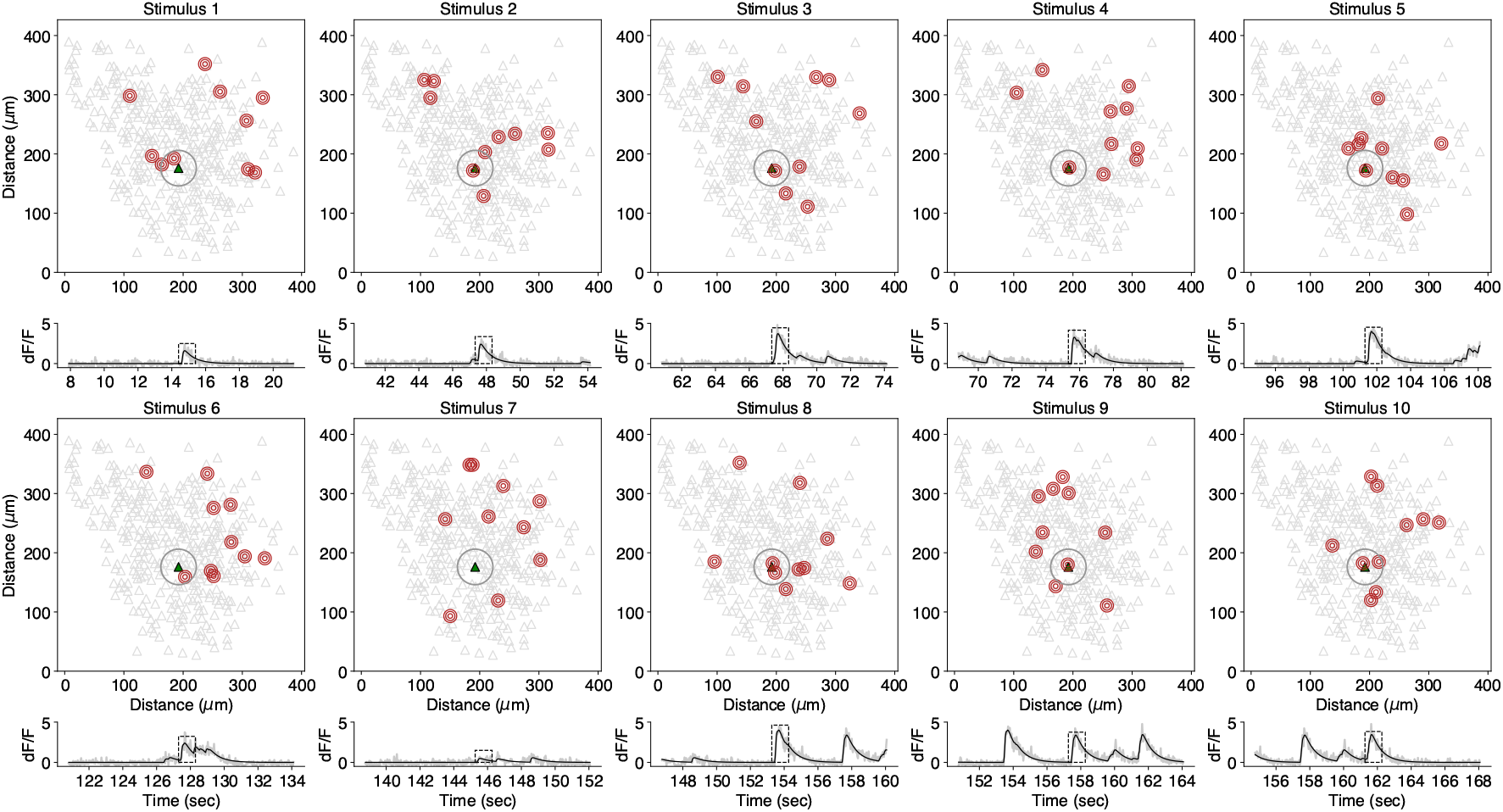
Experimental design and example neural responses during ORF mapping. Red circles represent simultaneously-delivered ensemble stimulation patterns (not drawn to scale). Green triangle enclosed by central gray circle represents an example neuron. Light gray traces below imaging FOV show raw dF/F signal elicited by the example neuron during stimulation; overlaid black traces show the denoised dF/F obtained using the OASIS algorithm. Dashed black rectangle shows the time-region where the stimulus-evoked response is calculated. Ten example stimuli are shown.

**Figure S2:**
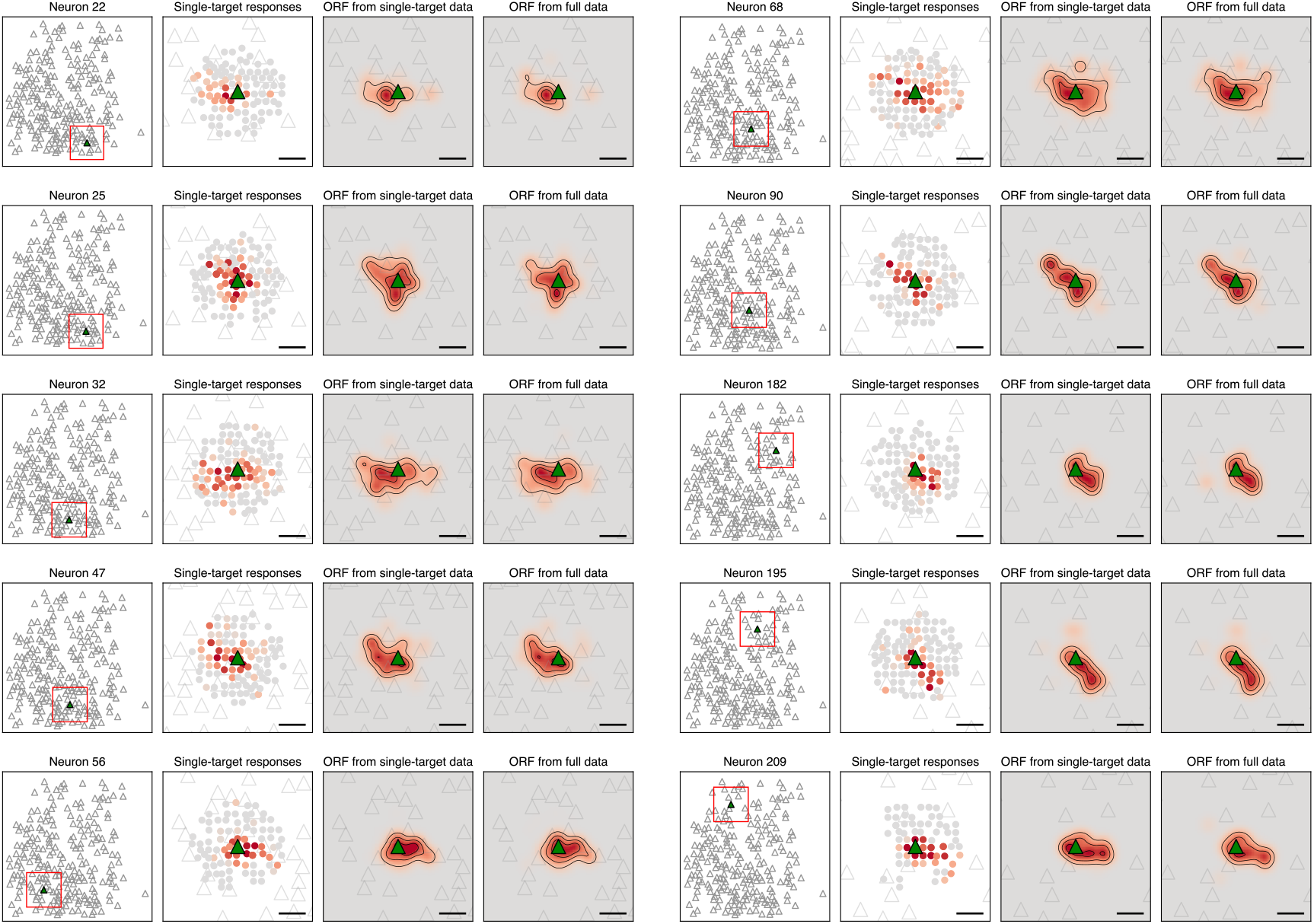
Sample optogenetic receptive fields from CA1 pyramidal neurons. Corresponds to the experiment in Figure 5. “Single-target responses” correspond to activity evoked by stimulation patterns when only one of ten hologram spirals was within 30 *µ*m of the example neuron, thus approximating ORFs that would be obtained by stimulating one location at a time. ORFs from single-target data are obtained by fitting the NBFR ORF model to such single-target responses. ORFs from full data are obtained similarly but using the complete dataset (not just fitting to the single-target responses). Scale bar, 20 *µ*m.

**Figure S3:**
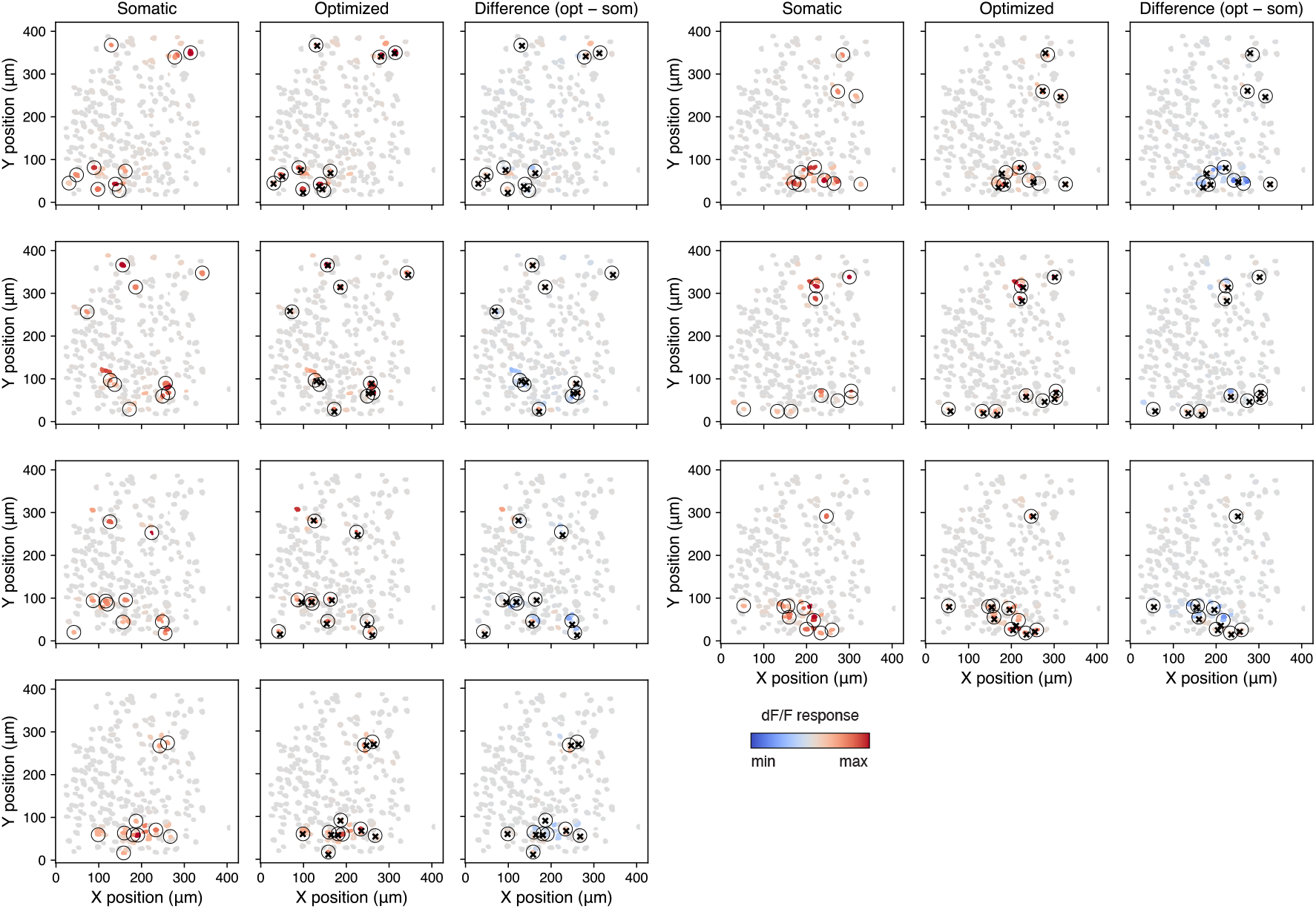
Evoked responses from all 7 ensembles in Figure 5. Black circles centered on target neurons. Optimized stimulus locations represented by black crosses. Responses are averages over 40 repetitions.

**Figure S4:**
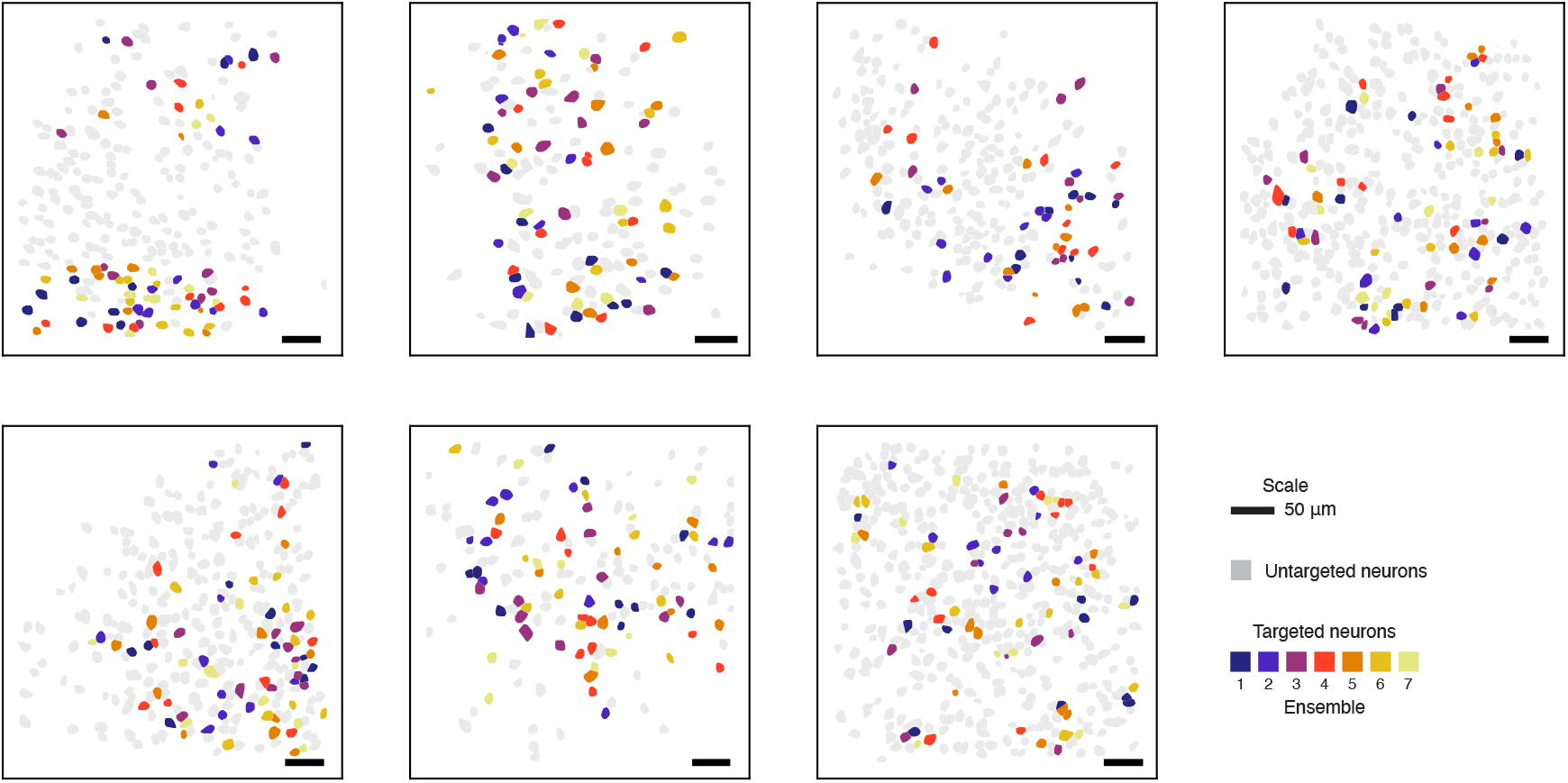
Tested ensembles from all 7 FOVs used in this study.

## References

[Abdeladim et al., 2023]Abdeladim, L., Shin, H., Jagadisan, U. K., Ogando, M. B., and Adesnik, H. (2023). Probing inter-areal computations with a cellular resolution two-photon holographic mesoscope. bioRxiv, pages 2023–03.

[Adesnik and Abdeladim, 2021]Adesnik, H. and Abdeladim, L. (2021). Probing neural codes with two-photon holographic optogenetics. Nature Neuroscience, 24(10):1356–1366.

[Antin et al., 2024]Antin, B., Sadahiro, M., Gajowa, M., Triplett, M. A., Adesnik, H., and Paninski, L. (2024). Removing direct photocurrent artifacts in optogenetic connectivity mapping data via constrained matrix factorization. PLOS Computational Biology, 20(5):e1012053.

[Baker et al., 2016]Baker, C. A., Elyada, Y. M., Parra, A., and Bolton, M. M. (2016). Cellular resolution circuit mapping with temporal-focused excitation of soma-targeted channelrhodopsin. Elife, 5:e14193.

[Bounds et al., 2023]Bounds, H. A., Sadahiro, M., Hendricks, W. D., Gajowa, M., Gopakumar, K., Quintana, D., Tasic, B., Daigle, T. L., Zeng, H., Oldenburg, I. A., et al. (2023). All-optical recreation of naturalistic neural activity with a multifunctional transgenic reporter mouse. Cell reports, 42(8).

[Buchanan et al., 2018]Buchanan, E. K., Kinsella, I., Zhou, D., Zhu, R., Zhou, P., Gerhard, F., Ferrante, J., Ma, Y., Kim, S. H., Shaik, M. A., et al. (2018). Penalized matrix decomposition for denoising, compression, and improved demixing of functional imaging data. BioRxiv, page 334706.

[Draelos et al., 2025]Draelos, A., Loring, M. D., Nikitchenko, M., Sriworarat, C., Gupta, P., Sprague, D. Y., Pnevmatikakis, E., Giovannucci, A., Benster, T., Deisseroth, K., et al. (2025). A software platform for real-time and adaptive neuroscience experiments. Nature communications, 16(1):9909.

[Draelos and Pearson, 2020]Draelos, A. and Pearson, J. (2020). Online neural connectivity estimation with noisy group testing. Advances in Neural Information Processing Systems, 33.

[Emiliani et al., 2022]Emiliani, V., Entcheva, E., Hedrich, R., Hegemann, P., Konrad, K. R., Lü scher, C., Mahn, M., Pan, Z.-H., Sims, R. R., Vierock, J., et al. (2022). Optogenetics for light control of biological systems. Nature Reviews Methods Primers, 2(1):55.

[Eybposh et al., 2020]Eybposh, M. H., Caira, N. W., Atisa, M., Chakravarthula, P., and Pégard, N. C. (2020). Deepcgh: 3d computer-generated holography using deep learning. Optics Express, 28(18):26636–26650.

[Giovannucci et al., 2019]Giovannucci, A., Friedrich, J., Gunn, P., Kalfon, J., Brown, B. L., Koay, S. A., Taxidis, J., Najafi, F., Gauthier, J. L., Zhou, P., et al. (2019). Caiman an open source tool for scalable calcium imaging data analysis. elife, 8:e38173.

[Grosenick et al., 2015]Grosenick, L., Marshel, J. H., and Deisseroth, K. (2015). Closed-loop and activity-guided optogenetic control. Neuron, 86(1):106–139.

[Hira, 2024]Hira, R. (2024). Closed-loop experiments and brain machine interfaces with multiphoton microscopy. Neurophotonics, 11(3):033405–033405.

[Hochbaum et al., 2014]Hochbaum, D. R., Zhao, Y., Farhi, S. L., Klapoetke, N., Werley, C. A., Kapoor, V., Zou, P., Kralj, J. M., Maclaurin, D., Smedemark-Margulies, N., et al. (2014). All-optical electrophysiology in mammalian neurons using engineered microbial rhodopsins. Nature methods, 11(8):825–833.

[Holmgren et al., 2003]Holmgren, C., Harkany, T., Svennenfors, B., and Zilberter, Y. (2003). Pyramidal cell communication within local networks in layer 2/3 of rat neocortex. The Journal of physiology, 551(1):139–153.

[Kaschube, 2014]Kaschube, M. (2014). Neural maps versus salt-and-pepper organization in visual cortex. Current opinion in neurobiology, 24:95–102.

[Knowles and Schwartzkroin, 1981]Knowles, W. D. and Schwartzkroin, P. A. (1981). Local circuit synaptic interactions in hippocampal brain slices. Journal of Neuroscience, 1(3):318–322.

[Lees et al., 2024]Lees, R. M., Pichler, B., and Packer, A. M. (2024). Contribution of optical resolution to the spatial precision of two-photon optogenetic photostimulation in vivo. Neurophotonics, 11(1):015006–015006.

[Mardinly et al., 2018]Mardinly, A. R., Oldenburg, I. A., Pégard, N. C., Sridharan, S., Lyall, E. H., Chesnov, K., Brohawn, S. G., Waller, L., and Adesnik, H. (2018). Precise multimodal optical control of neural ensemble activity. Nature neuroscience, 21(6):881–893.

[Marshel et al., 2019]Marshel, J. H., Kim, Y. S., Machado, T. A., Quirin, S., Benson, B., Kadmon, J., Raja, C., Chibukhchyan, A., Ramakrishnan, C., Inoue, M., et al. (2019). Cortical layer–specific critical dynamics triggering perception. Science, 365(6453):eaaw5202.

[Navarro and Oweiss, 2023]Navarro, P. and Oweiss, K. (2023). Compressive sensing of functional connectivity maps from patterned optogenetic stimulation of neuronal ensembles. Patterns.

[Pachitariu et al., 2016]Pachitariu, M., Stringer, C., Schröder, S., Dipoppa, M., Rossi, L. F., Carandini, M., and Harris, K. D. (2016). Suite2p: beyond 10,000 neurons with standard two-photon microscopy. BioRxiv, page 061507.

[Packer et al., 2012]Packer, A. M., Peterka, D. S., Hirtz, J. J., Prakash, R., Deisseroth, K., and Yuste, R. (2012). Twophoton optogenetics of dendritic spines and neural circuits. Nature methods, 9(12):1202–1205.

[Packer et al., 2015]Packer, A. M., Russell, L. E., Dalgleish, H. W., and Hä usser, M. (2015). Simultaneous all-optical manipulation and recording of neural circuit activity with cellular resolution in vivo. Nature methods, 12(2):140–146.

[Papagiakoumou et al., 2010]Papagiakoumou, E., Anselmi, F., Bégue, A., De Sars, V., Glückstad, J., Isacoff, E. Y., and Emiliani, V. (2010). Scanless two-photon excitation of channelrhodopsin-2. Nature methods, 7(10):848–854.

[Papagiakoumou et al., 2020]Papagiakoumou, E., Ronzitti, E., and Emiliani, V. (2020). Scanless two-photon excitation with temporal focusing. Nature Methods, 17(6):571–581.

[Pégard et al., 2017]Pégard, N. C., Mardinly, A. R., Oldenburg, I. A., Sridharan, S., Waller, L., and Adesnik, H. (2017). Three-dimensional scanless holographic optogenetics with temporal focusing (3d-shot). Nature communications, 8(1):1–14.

[Pnevmatikakis and Giovannucci, 2017]Pnevmatikakis, E. A. and Giovannucci, A. (2017). Normcorre: An online algorithm for piecewise rigid motion correction of calcium imaging data. Journal of neuroscience methods, 291:83–94.

[Prakash et al., 2012]Prakash, R., Yizhar, O., Grewe, B., Ramakrishnan, C., Wang, N., Goshen, I., Packer, A. M., Peterka, D. S., Yuste, R., Schnitzer, M. J., et al. (2012). Two-photon optogenetic toolbox for fast inhibition, excitation and bistable modulation. Nature methods, 9(12):1171–1179.

[Rickgauer and Tank, 2009]Rickgauer, J. P. and Tank, D. W. (2009). Two-photon excitation of channelrhodopsin-2 at saturation. Proceedings of the National Academy of Sciences, 106(35):15025–15030.

[Robinson et al., 2020]Robinson, N. T., Descamps, L. A., Russell, L. E., Buchholz, M. O., Bicknell, B. A., Antonov, G. K., Lau, J. Y., Nutbrown, R., Schmidt-Hieber, C., and Hä usser, M. (2020). Targeted activation of hippocampal place cells drives memory-guided spatial behavior. Cell, 183(6):1586–1599.

[Rolotti et al., 2022]Rolotti, S. V., Ahmed, M. S., Szoboszlay, M., Geiller, T., Negrean, A., Blockus, H., Gonzalez, K. C., Sparks, F. T., Canales, A. S. S., Tuttman, A. L., et al. (2022). Local feedback inhibition tightly controls rapid formation of hippocampal place fields. Neuron, 110(5):783–794.

[Seabold and Perktold, 2010]Seabold, S. and Perktold, J. (2010). Statsmodels: econometric and statistical modeling with python. SciPy, 7(1):92–96.

[Shemesh et al., 2017]Shemesh, O. A., Tanese, D., Zampini, V., Linghu, C., Piatkevich, K., Ronzitti, E., Papagiakoumou, E., Boyden, E. S., and Emiliani, V. (2017). Temporally precise single-cell-resolution optogenetics. Nature neuroscience, 20(12):1796–1806.

[Sridharan et al., 2022]Sridharan, S., Gajowa, M. A., Ogando, M. B., Jagadisan, U. K., Abdeladim, L., Sadahiro, M., Bounds, H. A., Hendricks, W. D., Turney, T. S., Tayler, I., et al. (2022). High-performance microbial opsins for spatially and temporally precise perturbations of large neuronal networks. Neuron.

[Thomson et al., 1996]Thomson, A. M., West, D. C., Hahn, J., and Deuchars, J. (1996). Single axon ipsps elicited in pyramidal cells by three classes of interneurones in slices of rat neocortex. The Journal of physiology, 496(1):81–102.

[Triplett et al., 2023]Triplett, M., Gajowa, M., Adesnik, H., and Paninski, L. (2023). Bayesian target optimisation for high-precision holographic optogenetics. Advances in Neural Information Processing Systems, 36:10972–10994.

[Triplett et al., 2025]Triplett, M. A., Gajowa, M., Antin, B., Sadahiro, M., Adesnik, H., and Paninski, L. (2025). Rapid learning of neural circuitry from holographic ensemble stimulation enabled by model-based compressed sensing. Nature Neuroscience, 28(10):2154–2165.

[Triplett and Goodhill, 2022]Triplett, M. A. and Goodhill, G. J. (2022). Inference of multiplicative factors underlying neural variability in calcium imaging data. Neural Computation, 34(5):1143–1169.

[Wagenmaker et al., 2024]Wagenmaker, A., Mi, L., Rozsa, M., Bull, M., Svoboda, K., Daie, K., Golub, M., and Jamieson, K. G. (2024). Active learning of neural population dynamics using two-photon holographic optogenetics. Advances in Neural Information Processing Systems, 37:31659–31687.

[Yang et al., 2018]Yang, W., Carrillo-Reid, L., Bando, Y., Peterka, D. S., and Yuste, R. (2018). Simultaneous two-photon imaging and two-photon optogenetics of cortical circuits in three dimensions. elife, 7:e32671.

[Zhang et al., 2018]Zhang, Z., Russell, L. E., Packer, A. M., Gauld, O. M., and Hä usser, M. (2018). Closed-loop alloptical interrogation of neural circuits in vivo. Nature methods, 15(12):1037–1040.

